# Reducing Cathodic Drift during Isoelectric Focusing using Microscale Immobilized pH Gradient Gels

**DOI:** 10.1101/2024.02.06.579215

**Authors:** Gabriela Lomeli, Amy E. Herr

## Abstract

Microfluidic analytical tools play an important role in miniaturizing proteomic assays for improved detection sensitivity, throughput, and automation. Microfluidic isoelectric focusing (IEF) can resolve proteoforms in lysate from low-to-single cell numbers. However, IEF assays often use carrier ampholytes (CAs) to establish a pH gradient for protein separation, presenting limitations like pH instability in the form of cathodic drift (migration of focused proteins toward the cathode). Immobilized pH gradient (IPG) gels reduce cathodic drift by covalently immobilizing the pH buffering components to a matrix. To our knowledge, efforts to implement IPG gels at the microscale have been limited to glass microdevices. To adapt IEF using IPGs to widely used microfluidic device materials, we introduce a polydimethylsiloxane (PDMS)-based microfluidic device and compare the microscale pH gradient stability of IEF established with IPGs, CAs, and a hybrid formulation of IPG gels and CAs (mixed-bed IEF). The PDMS-based IPG microfluidic device (μIPG) resolved analytes differing by 0.1 isoelectric point within a 3.5-mm separation lane over a 20-min focusing duration. During the 20-min duration, we observed markedly different cathodic drift velocities among the three formulations: 60.1 µm/min in CA-IEF, 2.5 µm/min in IPG-IEF (∼24-fold reduction versus CA-IEF), and 1.4 µm/min in mixed-bed IEF (∼43-fold reduction versus CA-IEF). Lastly, mixed-bed IEF in a PDMS device resolved green fluorescent protein (GFP) proteoforms from GFP-expressing human breast cancer cell lysate, thus establishing stability in lysate from complex biospecimens. µIPG is a promising and stable technique for studying proteoforms from small volumes.

---

Isoelectric focusing (IEF) is an equilibrium electrophoretic protein separation designed to resolve proteoforms, including proteins having an array of post-translational modifications (PTMs)^1,2^. During IEF, sample proteins electromigrate within a pH gradient and focus at the isoelectric point (pI) of each respective species. The pI is the pH at which a molecule has zero net charge, resulting in no electrophoretic mobility^3^. PTMs, such as phosphorylation and proteolytic cleavage, can alter the pI of a protein, making IEF an appropriate separation technique^4^. For instance, phosphorylation involves the addition of a negatively charged phosphate group to an amino acid, consequently reducing the pI of the protein^4^. The degree of change in pI is contingent upon factors like the number of PTM events, the specific amino acids involved, and the overall protein composition^4^.

Since the inception of IEF, miniaturization has been pursued as a means to achieve high-throughput analyses with reduced starting sample amounts, including for single-cell IEF analyses^2,5^. However, as IEF progresses towards the microscale, the increasing challenge of maintaining a stable pH gradient has limited the utility of this proteomic tool^6–8^. IEF can be achieved through various methods, such as using pH gradients generated via water electrolysis^9^ or thermal gradients^10^. Even with the introduction of novel methods for creating pH gradients, chemistry-based IEF technologies remain commonly used^11^. Here, we consider: carrier ampholyte IEF (CA-IEF)^12^, immobilized pH gradient IEF (IPG-IEF)^13^, and mixed-bed IEF (a hybrid of CA-IEF and IPG-IEF)^14–16^.

In CA-IEF, bracketed by anolyte (acidic) and catholyte (basic) regions and subjected to an applied electric field, a mixture of CAs arrange into an increasing pH gradient within an anticonvective matrix, such as polyacrylamide (PA) gel^17^. CAs are a mixture of small molecules with both positive and negative charge groups (amphoteric). CAs serve as a conductor of electrical current and act to buffer the pH^17^. CAs are generated with a chaotic synthesis method to generate hundreds of molecules capable of assembling into a monotonically increasing pH gradient^12^. Microfluidic IEF typically employs CA-IEF^2,7,18^. Unfortunately, CA-IEF suffers from pH gradient instability in the form of cathodic drift^19^. Cathodic drift is the observed movement of sample and other separation components toward the cathode, resulting in an overall shift in pH gradient^20^. The underlying mechanism of cathodic drift is attributed to several factors, including CA electromigration, electro-osmotic flow (EOF) due to the slight negative charge of PA gel, and electrolyte diffusion^21^. Cathodic drift leads to challenges during IEF, including loss of separation resolution^22^ and loss of proteins with high pH as the proteins run off the separation lane^7^. A prior microfluidic IEF study found >3-fold improvement in separation resolution when cathodic drift was reduced >20-fold^7^. While work has been done to reduce cathodic drift in centimeter-scale slab IEF to a manageable cathodic drift velocity (∼100 μm/min)^23^, cathodic drift velocity in microscale devices (∼10-600 μm/min)^18,21,24,25^ is more pronounced relative to the characteristic length-(micrometers) and timescales (min) for microscale IEF separation^6^.

IPG-IEF was developed to improve pH gradient stability over CA-IEF^19^. In IPG-IEF, a pH gradient is established prior to sample application by copolymerizing a class of buffering molecules known as Immobilines into PA gel. By creating a concentration gradient of Immobilines at the time of polymerization, an immobilized pH gradient (IPG) gel is established. Unlike CAs, Immobilines are not amphoteric molecules, but are instead either weak acid or weak base acryloyl monomers that can be polymerized into the PA gel to participate in protolytic equilibria to perform their buffering function^26^. Cathodic drift is reduced or even eliminated in IPG-IEF, because the Immobilines are polymerized into the PA gel, and therefore insolubilized, thus the resultant pH gradient remains fixed in space and does not drift under the action of an applied electric field^26,27^. A microfluidic IPG device was previously developed in glass microchannels^28^. While a cornerstone of early microfluidic advances—glass microfluidic devices have seen commercial success^29^ and continue to be utilized—a next wave of microfluidic researchers are moving from glass-based devices to the rapid prototyping possible with polymer-based devices^30,31^.

While useful for mitigating cathodic drift, IPG-IEF does require longer focusing times^16,32^ and higher sample amounts^16^ versus CA-IEF, so another IEF technology, mixed-bed IEF, was introduced to combine the advantages of both IPG- and CA-IEF^19^. In mixed-bed IEF, a stationary, Immobiline-driven pH gradient coexists with a soluble, CA-driven pH gradient^15^. The Immobiline molecules are hypothesized to suppress EOF and stabilize the CA-driven pH gradient^33^, similar to how prior work has shown that the incorporation of a tertiary base in PA gel, in stoichiometrically matched amounts compared to negative charges, can reduce cathodic drift due to EOF^23^. Mixed-bed IEF recovers the focusing speed and sensitivity of CA-IEF^16^, while retaining the pH stability of IPG-IEF^15^. Importantly, prior efforts to implement microscale mixed-bed IEF have been unsuccessful for reasons that were not elaborated^28^, a question explored in the present study.

In this work, we consider polydimethylsiloxane (PDMS) microdevices, hereafter referred to as µIPG devices, as platforms for IPG-IEF and mixed-bed IEF. We explore PDMS as a separations-device substrate owing to its cost-effective, rapid prototyping potential and biocompatibility^34^. First, we devise a protocol to fabricate IPG gels in PDMS micro-channels borrowing benzophenone-assisted PA gel polymerization from our earlier reported work^31^. We next investigate mitigation of cathodic drift in the μIPG devices when operated with CA-IEF, IPG-IEF, or mixed-bed IEF modes. Finally, we perform mixed-bed IEF in the µIPG device for analysis of green fluorescent protein (GFP) proteoforms from crude mammalian cell lysate.

## EXPERIMENTAL SECTION

### µIPG Device Design

The μIPG device was fabricated by standard soft lithography methods. The PDMS substrate was patterned with 4 microchannels and assembled onto a glass slide. Each microchannel is 3.5-mm long, 100-μm wide, and 50-μm tall. We apply Kapton tape on the glass slide opposite of the PDMS substrate to serve as a photomask and prevent photopolymerization in those regions. A detailed protocol is described in the Supporting Information.

### Polyacrylamide Gel Fabrication

To polymerize PA gel in the PDMS channels, we adapted a technique from previous work which ensures gel polymerization in a microchannel environment that may be O_2_ rich^31^. Briefly, the microchannels are treated with benzophenone before any gel precursor is applied. Benzophenone serves as an oxygen scavenger when exposed to UV^35^. Next, 6 %T PA gel precursor is applied at one of the microchannel inlets. Supplemental Table S1 summarizes the PA gel precursor recipes. The scaffold PA gel precursor is then photopolymerized with UV light. The UV activates both the benzophenone (oxygen scavenger) and the photoinitiator in the PA gel precursor (radical polymerization initiator).

The inlets were then emptied and filled with 6 %T or 12 %T PA acidic (pH = 3.8) and basic (pH = 7.0) Immobiline-PA gel precursor solutions. Acidic Immobilines used were acrylamido buffers pKa 3.6 and pKa 4.6. Basic Immobilines used were acrylamido buffers pKa 6.2, pKa 7.0, and pKa 9.3. The acidic and basic IPG precursor recipes were adapted from previous work^28^. The precursors were allowed to diffuse into the micro-channels for 7 hr to establish a linear concentration gradient along the length of the microchannel. During this diffusion step, the polymerized 6 %T PA gel serves as a scaffold for the IPG gel, maintaining quiescent conditions during gel precursor diffusion^25^. The IPG precursor was then photopolymerized with UV light. The devices were incubated with (for CA-IEF or mixed-bed IEF) or without (for IPG-IEF) 1% ZOOM Carrier Ampholytes pH 4 – 7 dissolved in DI water or sample loading buffer overnight (as summarized in Supplemental Table S2). Detailed PA gel precursor reagents, photopolymerization conditions, and the composition of the sample loading buffer are described in the Supporting Information. Modifications to the PA gel fabrication protocol for pH gradient characterization and polymerization experiments are also detailed in the Supporting Information.

### Cell Culture and Cell Lysate Preparation

An MCF7 human breast cancer cell line genetically modified to stably express enhanced green fluorescent protein (GFP) was obtained from the American Type Culture Collection and maintained using standard cell culture practices. Cell lysate preparation was performed as previously described^18^ with some modifications. A detailed procedure is described in the Supporting Information.

### Isoelectric Focusing Experiments

Fluorescent pI markers (pIs 4.5, 5.5, 5.9, 6.6, and 6.7) were used at various concentrations to provide similar intensities. For cell lysate experiments, 1 μL of MCF7-GFP cell lysate (∼20,000 cells/µL) was applied to both the anode and cathode inlets (∼40,000 cells total), as well as pI markers in sample loading buffer. Supplemental Table S3 lists the anode inlet and cathode inlet sample components for all IEF experiments. An electric field was applied using the following voltage ramp: 50 V/cm for 4 min, 100 V/cm for 5 min, 200 V/cm for 5 min, and 300 V/cm for 6 min or more depending on the experiment. The detailed IEF procedure is described in the Supporting Information.

### Imaging and Image Analysis

Microscope information for the fluorescence imaging as well as the methods for image/micrograph analysis are described in the Supporting Information.

## RESULTS AND DISCUSSION

### Design of μIPG for Isoelectric Focusing

The IEF device can be operated using CA-IEF, IPG-IEF, or mixed bed IEF (Figure 1A). The IEF device comprises 4 separation lanes within the footprint of a standard microscope slide (Figure 1B). During IEF, loaded analytes migrate along the pH gradient within the separation lane until each analyte reaches the location in the pH gradient where net zero charge arises in the molecule (Figure 1C).

**Figure 1.**
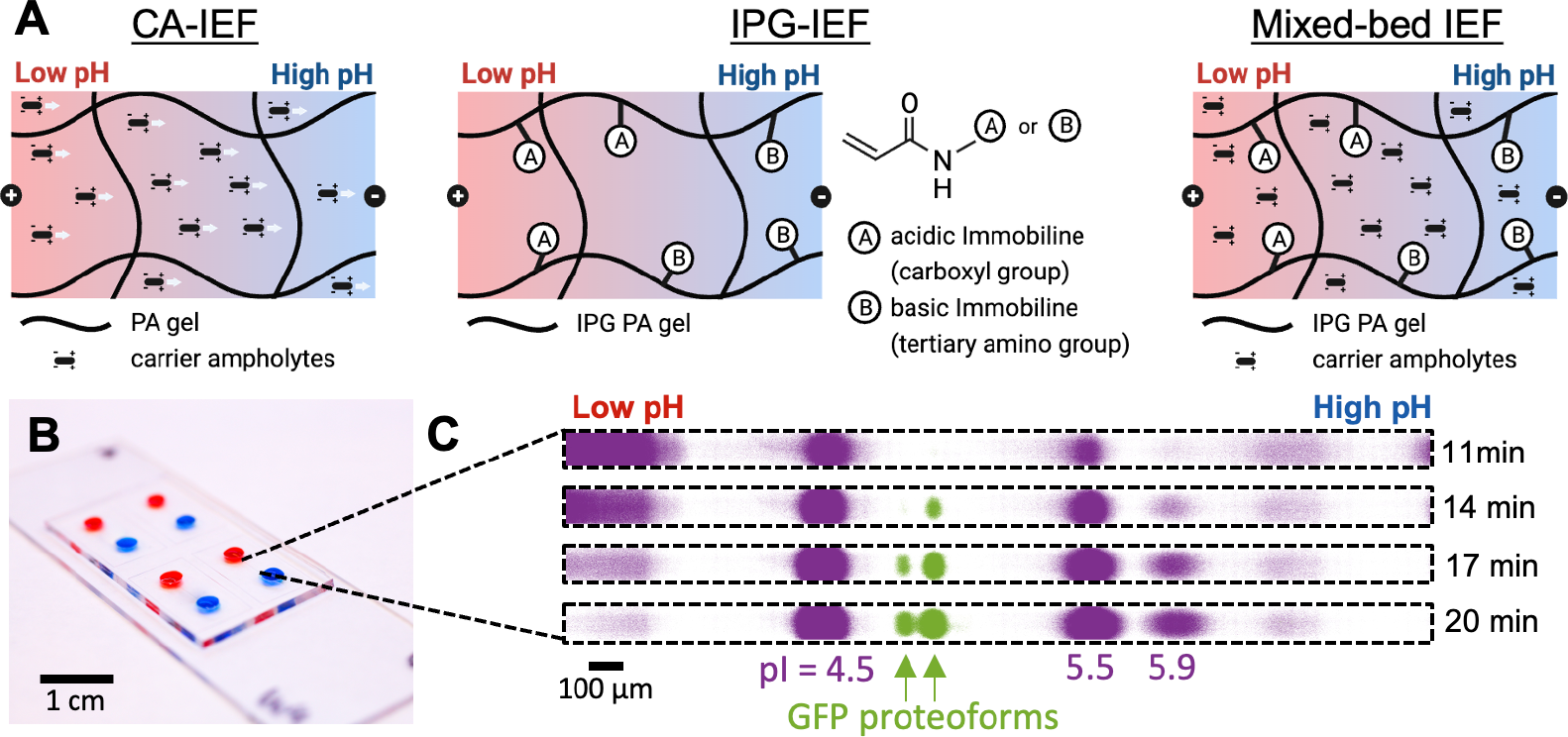
µIPG is designed to mitigate cathodic drift during IEF at the microscale for stable pI-based separation of proteoforms. (A) Schematic of pH buffering mechanisms for CA-IEF, IPG-IEF, and mixed-bed IEF. In CA-IEF, the pH is buffered by carrier ampholytes, which are subject to cathodic drift (indicated by gray arrows). In IPG-IEF, Immobilines buffer the pH and cathodic drift is mitigated by covalent attachment of Immobilines to the PA gel. In mixed-bed IEF, the Immobilines stabilize the CA pH gradient to reduce cathodic drift. (B) µIPG device containing food dye-filled microchannels. (C) Inverted fluorescence micrographs of mixed-bed IEF-separated pI markers (pIs 4.5, 5.5, and 5.9) and two GFP proteoforms at several time points demonstrates pH gradient stability as pI marker and protein peaks remain fixed in position over time. Micrographs have the same acquisition settings, brightness, and contrast. Representative of n = 3 separations. IPG gel is 6 %T.

We first sought out to validate IPG-IEF in our IEF device before incorporating CAs for mixed-bed IEF. To fabricate a scaffold and IPG gel, Figure 2A reports a double-photopoly-merization process. Steps i-v establish a scaffold gel within the PDMS microchannel, while steps vi-ix overlay the IPG onto the PA scaffold through diffusion, creating a composite hydrogel/interpenetrating network. The scaffold gel prevents fluid flow and enables the establishment of a linear Immobiline gradient in steps vii. A similar strategy was previously employed to create an on-chip pore-size gradient in PA gel^25^. UV irradiation is used in both photopolymerization steps for two reasons. Firstly, the UV activates the benzophenone previously absorbed into the PDMS to quench oxygen and graft the PA gel to the PDMS^35^. Secondly, the UV activates the photoinitiator, VA-086, in the gel precursors to initiate radical PA polymerization. We confirmed that all precursors polymerized in the PDMS channel and that both UV irradiation and benzophenone were necessary for polymerization (Supplemental Figure S1). Therefore, we overcome the challenge of performing radical chemistry in PDMS by utilizing an oxygen scavenger^35^. Finally, the choice of buffer in step ix dictates whether the device is operated as IPG-IEF (no CAs in the buffer) or mixed-bed IEF (CAs included in the buffer).

**Figure 2.**
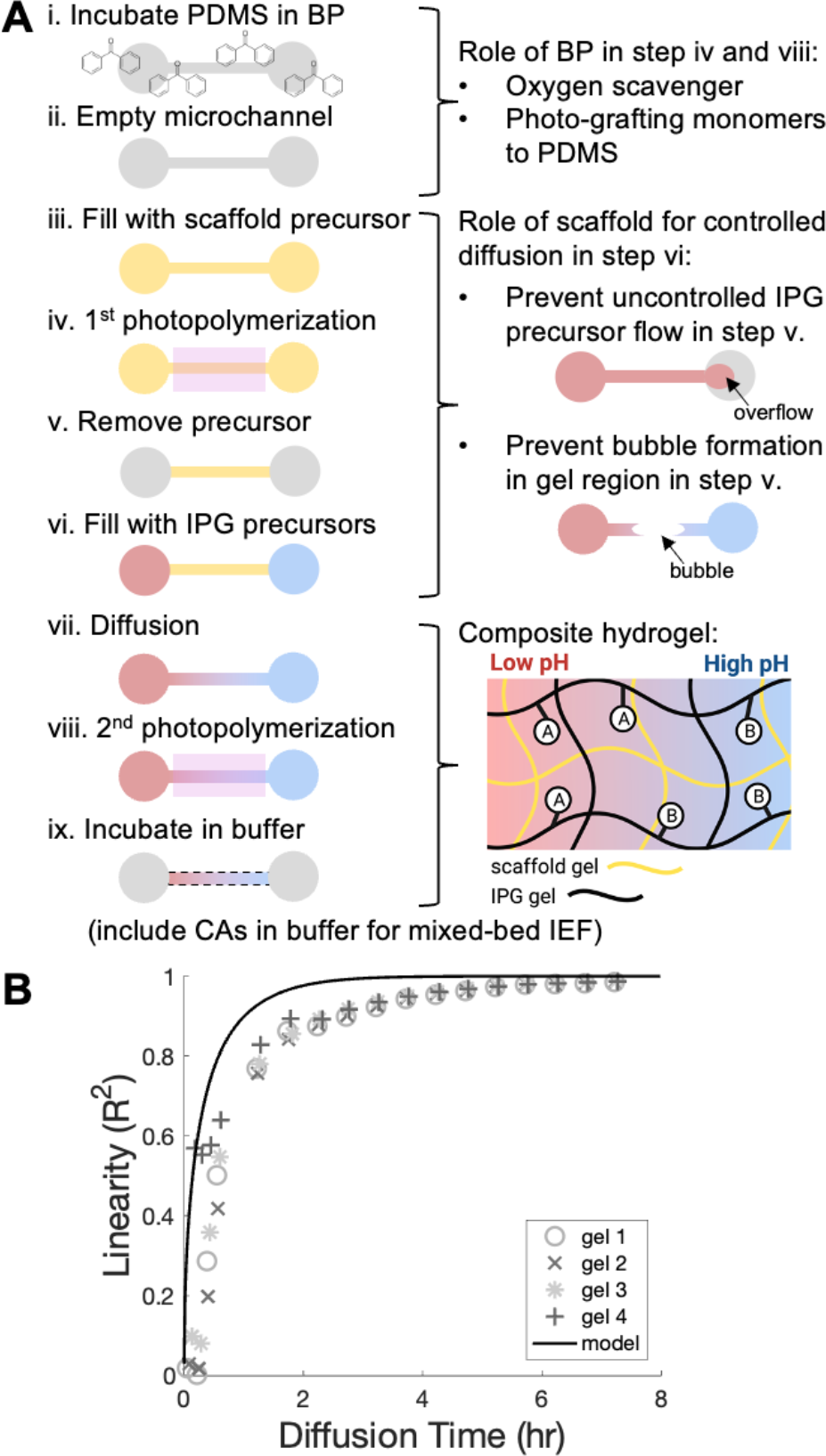
Design and fabrication of the µIPG device. (A) µIPG device fabrication protocol using a double photopolymerization method. (B) Linearity of gradient during diffusion step is modeled and experimentally monitored (n = 4 gels) with Nile Blue acrylamide dye.

### Establishing a Linear pH Gradient

We sought to establish a linear Immobiline concentration gradient within the μIPG microchannel, which in turn creates a linear pH gradient. IPG pH gradients are highly tunable, including the option to form a linear or nonlinear gradient^19^. A nonlinear pH gradient may be desired to create regions of narrow pH to analyze complex protein mixtures^36,37^. To demonstrate the feasibility of IPG-IEF in a PDMS device, the present work focuses on linear pH gradients.

To determine the diffusion time necessary to establish a linear gradient of IPG gel precursor molecules within a microchannel prefilled with 6 %T PA scaffold gel, we modeled the diffusion behavior of a proxy molecule, Nile Blue acrylamide, that is detectable using real-time fluorescence imaging (unlike Immobiline molecules). Since Nile Blue acrylamide has a similar, yet larger, molecular mass than all the individual molecules of the IPG gel precursors, we can therefore assume Nile Blue acrylamide will have a similar or slower diffusion rate than the IPG gel precursor molecules we want to model. The diffusion coefficient for Nile Blue acrylamide is calculated to be 0.0162 mm^2^/min (Supplemental Note S1).

Next, the concentration profile of Nile Blue acrylamide in the microchannel was modeled with the 1-D diffusion equation (Supplemental Note S1). We compared the numerical modeling results to experimental results and observed that diffusion in the microchannel was slower than predicted (R^2^ > 0.95 achieved after ∼250 min instead of expected ∼120 min) (Figure 2B). We hypothesize that the slower effective Nile Blue acrylamide diffusion could be due to absorption of Nile Blue acrylamide by the PDMS^38^, which we observed in our devices (data not shown). Based on our experimental results, the Nile Blue acrylamide concentration profile is linear after 7 hr (Figure 2B). We therefore chose a diffusion time of 7 hr as a conservative estimate of time needed to establish a linear Immobiline gradient, which allowed device fabrication to be completed in one day.

After fabricating µIPG devices with Immobiline reagents, we next sought to investigate the linearity of the pH gradient via IEF of a fluorescent pI marker ladder. Our μIPG device was designed to separate analytes having pI’s in the pH range of 3.8 to 7.0. Figure 3A demonstrates separation of 5 pI markers (ranging from 4.5 to 6.7). The pH gradient in the µIPG device is linear from 4.5 to 6.7 (Figure 3B). We observe device-to-device variation in the position of the pH gradient within the microchannel (i.e., gel 2 in Figure 3B). We hypothesize that this variation was caused by the visual alignment of the Kapton tape photomask to the microchannels during device fabrication. Reduction in device-to-device variability could be achieved using a mask aligner. To account for the relative local pH conditions, we include fluorescent pI markers in all IEF runs to assess run success and as internal standards of pI, as is routine in IEF^39^.

**Figure 3.**
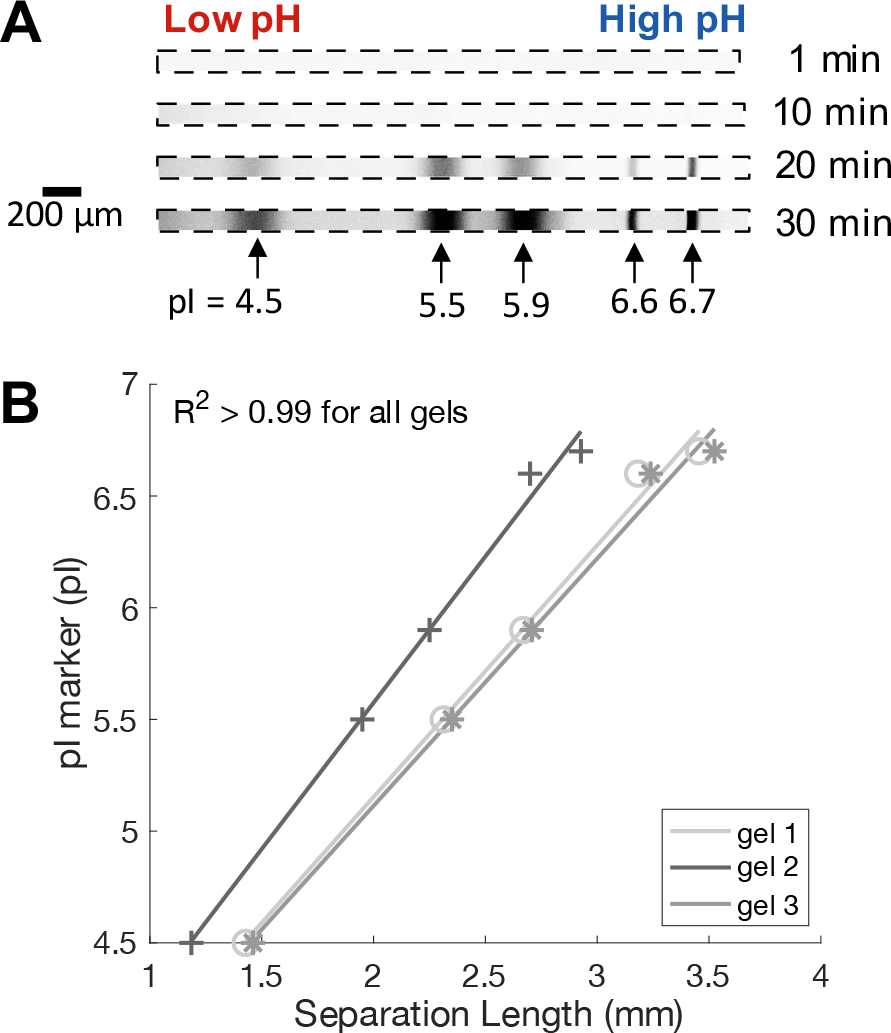
IEF of pI markers in µIPG device confirms a linear pH gradient. (A) Inverted fluorescent micrographs of IEF-separated pI markers at several time points demonstrates separation evolution and stability. Micrographs have the same acquisition settings, brightness, and contrast. Representative of n = 3 separations. (B) Plot shows the 5 pI markers versus position to determine the linearity of the pH gradient from pH 4.5 – 6.7 for n = 3 gels/separations. IPG gel

Notably, the μIPG device resolves pI markers 6.6 and 6.7, which differ by 0.1 pH unit (Figure 3). For context, a single phosphorylation event can cause a pI change of 0.3-0.4 pH units^40^. We anticipate that even smaller pI differences can be resolved in the μIPG device for proteins (∼100-fold larger molecular mass than pI markers), since smaller molecular mass samples tend to experience more peak broadening and therefore loss in separation resolution^41^. Moreover, the pH gradient in IPGs is highly tunable, so increased separation resolution may also be achieved by choosing a narrower pH range than the 3.8-7.0 pH range employed here. Critically, our PDMS-based μIPG device performance in terms of minimum resolvable pI difference (Δ(pl)_min_∼0.1) is comparable to the previous glass-based μIPG device (Δ(pl)_min_∼0.04)^28^, even though the separation length in the PDMS-based μIPG device is 1.7-fold shorter (3.5 mm versus 6 mm), making our PDMS-based µIPG device the μIPG device with the shortest separation length to our knowledge. In addition to reducing the footprint of each separation, which can have advantages for increasing throughput, we anticipate that additional miniaturization of the μIPG device will facilitate analysis of small sample amounts by reducing the available surface area for nonspecific absorption of sample to the microchannel walls and anticonvective matrix. Moreover, since the peak positions are stationary (Figure 3A), the separation lane of the µIPG device can remain short compared to CA-based microfluidic devices where the microchannel should be sufficiently long to prevent samples from running off the separation lane due to cathodic drift^7^.

### Stability and Performance of µIPG Device

To compare pH gradient stability in CA-IEF, IPG-IEF, and mixed-bed IEF, we monitored pI marker peak positions over time in our μIPG device for all three conditions. To perform an accurate comparison between IPG-IEF and CA-IEF in our device, the only modifications we made to the CA-IEF condition compared to the IPG-IEF condition are: (1) Immobiline reagents were excluded from the PA gel, and (2) the device was incubated in 1% CAs in DI water overnight. The mixed-bed IEF condition included Immobiline reagents and was incubated in 1% CAs in DI water overnight. Anode and cathode sample components were the same for all IEF conditions (Supplemental Table S2).

Figure 4A kymographs demonstrate a representative separation evolution for all three IEF conditions tested and Figure 4B summarizes the peak drift behavior for pI marker 5.5 for all conditions. Based on the data collected during 20 min of IEF (Figure 4A-B, Supplemental Figure S2), the cathodic drift velocity of pI marker 5.5 is 60.1 ± 7.7 µm/min for CA-IEF, 2.5 ± 2.5 µm/min for IPG-IEF (∼24-fold reduction versus CA-IEF), and 1.4 ± 0.4 µm/min for mixed-bed IEF (∼43-fold reduction versus CA-IEF). Interestingly, we also observe anodic drift for pI marker 4.5 in one of the CA-IEF conditions (Supplemental Figure S2), which is another source of pH gradient instability^20^. Additionally, CA-IEF in our device did not resolve pI marker 5.9. We hypothesize that cathodic drift caused pI marker 5.9 to “run off” the gel before pI marker 5.9 was sufficiently concentrated to be detected. Our µIPG device, operated with either IPG-IEF (2.5 μm/min cathodic drift velocity) or mixed-bed IEF (1.4 μm/min cathodic drift velocity), represents a marked improvement in pH gradient stability compared to CA-IEF (60.1 μm/min cathodic drift velocity) and previously published microscale CA-IEF devices (∼10-600 μm/min cathodic drift velocities)^18,21,24,25^.

**Figure 4.**
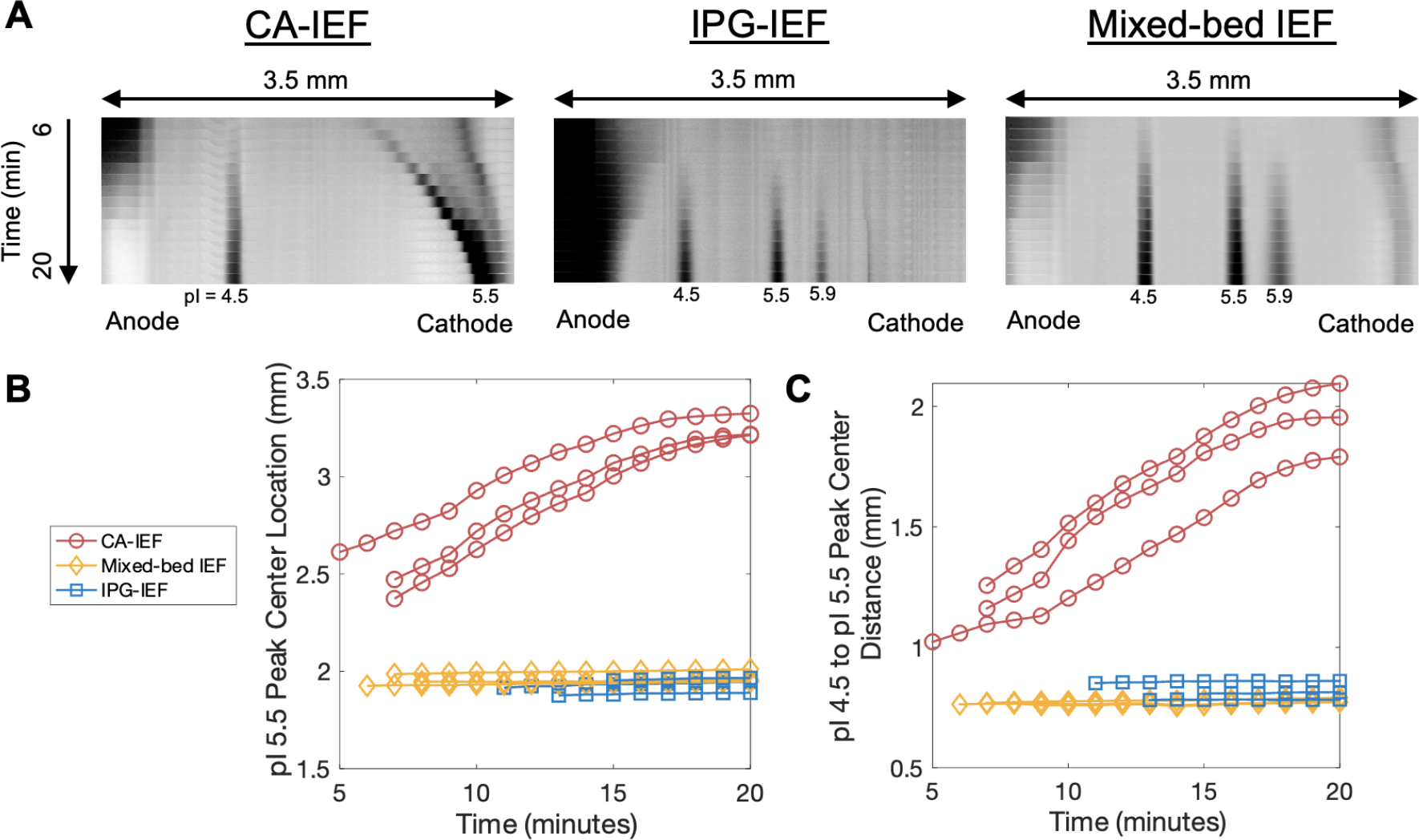
Cathodic drift in µIPG is reduced with IPG- or mixed-bed IEF versus CA-IEF. (A) Inverted fluorescence kymographs of IEF-separated pI markers demonstrate separation evolution. (B) Plot of pI 5.5 peak center location versus IEF focusing time shows that the peak center has a stable position for IPG-IEF and mixed-bed IEF (minimal cathodic drift), while the peak center drifts towards the cathode for the CA-IEF separations. (C) Plot of distance between pI 4.5 and pI 5.5 peak centers versus IEF focusing time demonstrates pH gradient shape is more stable for IPG- and mixed-bed IEF versus CA-IEF. For (B) and (C), peak center data from n = 3 separations for each IEF condition are plotted for each time point with SNR > 3.

Next, we sought to assess whether the pH gradient compressed or expanded over the course of IEF for the three IEF configurations scrutinized. Since the CA-IEF condition lacked the third pI marker (pI marker 5.9) needed to measure the slope of the pH gradient (dpH/d*x*) as would assess pH gradient shape, we instead measured the distance between pI markers 4.5 and 5.5 over time as a proxy for pH gradient shape (Figure 4C). The distance between pI markers 4.5 and 5.5 remained stable for IPG-IEF and mixed-bed IEF, while the CA-IEF condition saw an ∼2-fold increase in the distance between pI markers 4.5 and 5.5 during 20 min of IEF (Figure 4C). Additionally, the positions of 5 pI markers between 4.5 and 6.7 remained stable over the course of 30 min IEF using IPG-IEF (Figure 3A). As expected of IPG-based IEF separations, where pH-generating species are covalently fixed in the anticonvective matrix^26^, pI marker peak positions for both IPG-IEF and mixed-bed IEF remained stable compared to the CA-IEF condition.

Lastly, we evaluated the focusing performance of the µIPG device. Using the relative analyte concentration within the focused peaks at several time points during IEF, we can assess the assay sensitivity of the three IEF conditions (i.e., higher analyte concentrations correlate with higher assay sensitivity). The AUC (a proxy for analyte concentration) in arbitrary fluorescence units for pI marker 4.5 at 20 min in our µIPG device is 8697 ± 4568 for CA-IEF (which we will define as max AUC), 5547 ± 1568 for mixed-bed IEF (64% of max AUC), and 1735 ± 1232 for IPG-IEF (20% of max AUC) (Figure 5A, n = 3 separations). Additionally, CA-IEF and mixed-bed IEF reached SNR > 3 faster than IPG-IEF (Figure 5B). While IPG-IEF has a more stable pH gradient than CA-IEF (Figure 4), IPG-IEF has poorer focusing performance than CA-IEF with regards to detection sensitivity and focusing time (Figure 5). Mixed-bed IEF has improved focusing performance versus IPG-IEF alone (Figure 5), while retaining pH gradient stability (Figure 4). Differences between the three IEF conditions in AUC at 20 min (Figure 5A) and the time to reach SNR > 3 (Figure 5B) are not statistically significant using a One-way ANOVA with Kruskal−Wallis test and post-hoc Dunn’s multiple comparison test (*p* < 0.05). However, based on the average results of n = 3 separations, our microscale experiments seem to corroborate prior findings that mixed-bed IEF improves detection sensitivity and reduces the focusing time versus IPG-IEF alone^16^.

**Figure 5.**
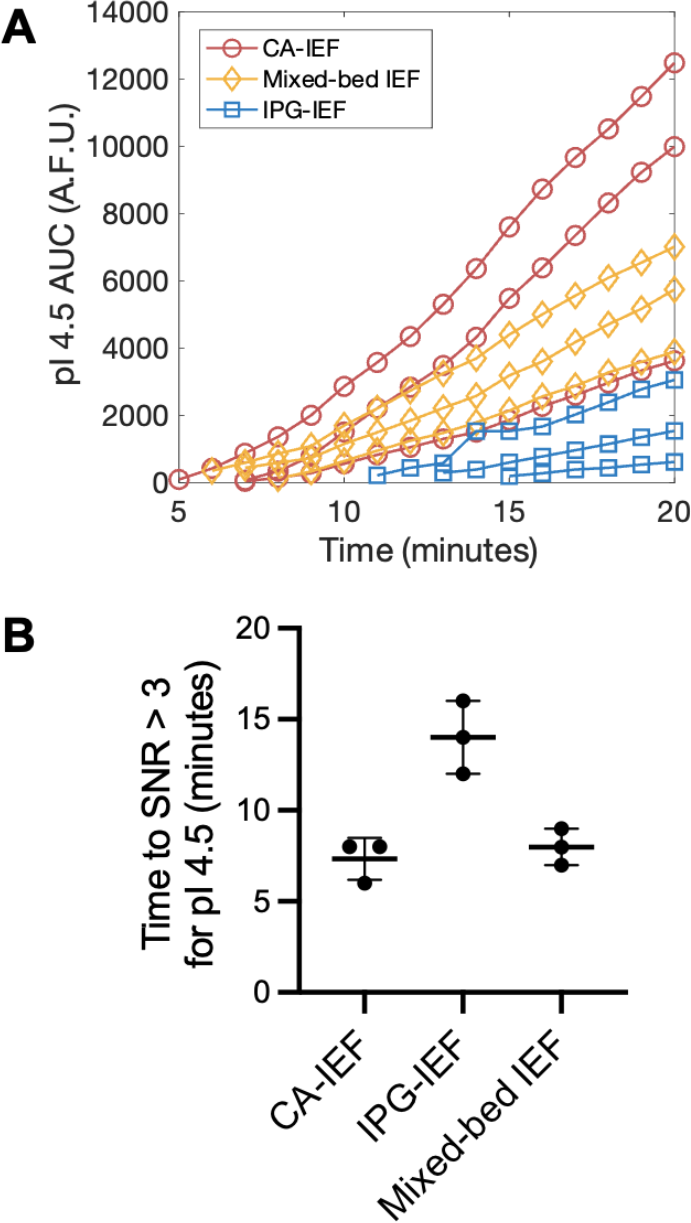
Comparing the sensitivity and focusing time of IPG-IEF, CA-IEF, and mixed-bed IEF in a microfluidic IEF device. (A) Plot of pI 4.5 marker AUC versus IEF focusing time shows that mixed-bed IEF improves pI marker concentration versus IPG-IEF alone. AUC data from n = 3 separations for each IEF condition are plotted for each time point with SNR > 3. (B) Time to SNR > 3 for 3 IEF conditions demonstrates relative focusing speeds.

### Cell Lysate Separations

To understand the performance of the μIPG device with a complex protein sample, we performed IEF of MCF7-GFP cell lysate. IEF was performed using both a 6 %T IPG gel formulation (Figure 1C) and a 12 %T IPG gel formulation (Figure 6). Cell lysate IEF experiments were first performed using IPG-IEF, yet IPG-only IEF did not successfully resolve the pI markers or GFP proteoforms (Figure 6). We hypothesized that the salts in the cell lysate buffer could be interfering with IEF in the IPG-only device^42^, so we tested mixed-bed IEF in the μIPG device (Figure 6). Mixed-bed IEF is beneficial in samples containing high salt levels where the presence of CA in the sample improves buffering^42^. Mixed-bed IEF is achieved by the addition of 1% CAs to the IPG gel, which we accomplished by incubating the μIPG device in a sample loading buffer containing 1% CAs overnight.

**Figure 6.**
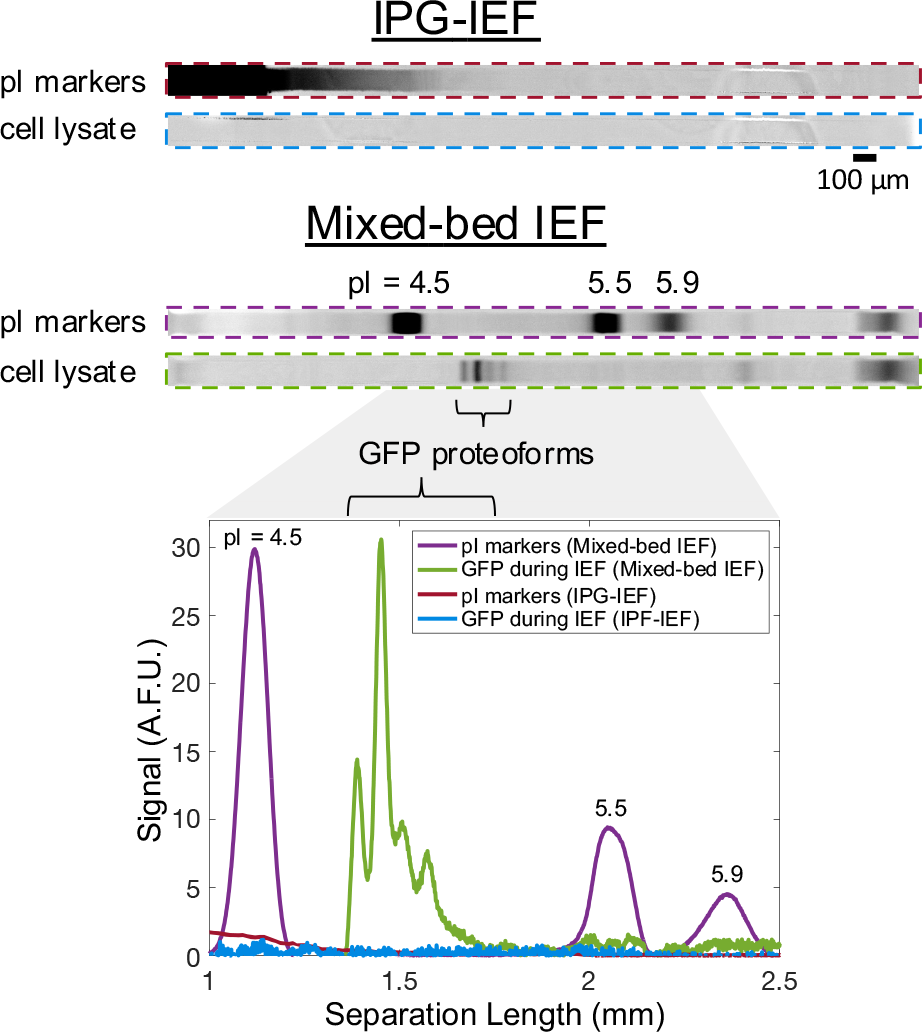
Mixed-bed IEF resolves pI markers and GFP proteoforms from MCF7-GFP cell lysate. Inverted fluorescence micrographs and corresponding intensity plot of pI markers and GFP focused in µIPG device from MCF7-GFP cell lysate (40,000 cells). IEF performed using IPG-IEF or mixed-bed IEF. Profiles captured after 20 min of IEF. IPG gel is 12 %T. Micrographs in the same channel have the same acquisition settings, brightness, and contrast. Representative of n = 3 separations for IPG-IEF and n = 14 separations for mixed-bed IEF.

IEF of MCF7-GFP cell lysate in the μIPG device using mixed-bed CA-IEF resolved up to four GFP proteoforms (Figure 6). Out of 14 independent IEF separations (technical replicates), 1 separation detected only one GFP proteoform, 6 separations resolved two GFP proteoforms, 3 separations resolved three GFP proteoforms, and 4 separations resolved four GFP proteoforms (Supplemental Figure S3). Variation in the number of proteoforms resolved could be attributable to the manual fabrication of μIPG devices, as discussed in the *Performance of μIPG device* subsection. Additionally, variation in the distance between electrodes during μIPG operation could also lead to variability in electric field strengths between technical replicates. The large size of the reservoirs (3-mm diameter) meant that the distance between the electrodes could vary from 3.5 mm to 9.5 mm, and therefore the final electric field strength could vary from 284 V/cm to 771 V/cm. Since separation resolution is dependent on electric field strength^17^, changes in electric field strength between different separations could lead to peak broadening that would obscure proteoform peaks. Mechanically fixing the distance between electrodes could be an avenue to better control electric field strength, and therefore, separation resolution. The GFP proteoform data (Figure 6, Supplemental Figure S3) serve as proof-of-concept that a complex protein sample can be analyzed in a µIPG device with mixed-bed IEF without introducing additional complexity (CAs were loaded diffusively instead of electrophoretically).

The GFP proteoforms detected had pIs between ∼4.8 and 5.0, comparable to the glass-based μIPG device (GFP pIs ranging from 4.88 to 5.19^28^) and slab-gel IEF (GFP pIs ranging from 4.7 to 5.1^43^). The various GFP proteoforms have been attributed to differential C-terminal cleavage by non-specific proteases^43^. While the core of the GFP protein is resistant to proteolysis, the C-terminus “tail” sequence, His-Gly-Met-Asp-Glu-Tyr-Lys, contains both basic and acidic amino acid residues, which, when cleaved, produce a variety of GFP proteoforms^43^. These GFP proteoforms can be detected by various techniques, including isoelectric focusing^44^ and capillary zone electrophoresis^45,46^.

## CONCLUSIONS

The design and development of IEF using IPG gels in elastomeric microfluidic devices advances microfluidic analytical tools for proteomic assays by allowing rapid prototyping device design approaches to be adopted for protein separations. The double photopolymerization procedure reported here for fabrication of μIPG creates a composite gel, which facilitates IEF through the IPG gel component, allowing for resolution of analytes differing by approximately 0.1 isoelectric point. Motivated by the desire to design for enhanced IEF assay throughput, minimizing the separation channel length allows for more separation lanes per unit surface area. The substantial cathodic drift observed in microscale IEF has limited designing to these performance goals to date. Recognizing the cathodic drift limitation on microfluidic IEF, we demonstrate that the µIPG device operated using mixed-bed IEF reduces cathodic drift compared to CAIEF. Further, the mixed-bed IEF configuration allows for analysis of cell lysate not previously reported with microscale IPG-IEF alone. By addressing the limitations of carrier ampholytes and slow-prototype-cycle glass devices, the μIPG device described here offers a promising platform for protein separation and analysis.

Future research can focus on expanding the capabilities of the μIPG by exploring different gel formulations (i.e., to modify the pH gradient). Additionally, efforts can be directed towards enhancing the sensitivity and resolution of the μIPG for the analysis of complex protein samples, enabling the detection and characterization of rare or low abundance proteoforms. Moreover, the flexibility of PDMS allows for the creation of complex microfluidic designs and the integration of various features, such as channels, valves, and chambers, in a single device. We anticipate that the PDMS-based μIPG device presented here could be integrated with other PDMS-based microfluidic devices for additional capabilities (i.e., to automate sample preparation). Lastly, the reversible bond between PDMS and glass could be used to expose the IPG gel for immunostaining of proteins^31^.

With an ability to analyze complex biological samples and achieve high-resolution protein separations, the μIPG device holds great potential for advancing the field of proteomics and undergirding further miniaturization of analytical techniques.

## ASSOCIATED CONTENT

### Supporting Information

Supplemental Experimental Section; Both benzophenone coating and UV irradiation are necessary for PA gel polymerization in the PDMS device; Intensity plots and micrographs of pI markers 4.5, 5.5, and 5.9 focused in microfluidic IEF devices in CA-IEF, IPG-IEF, and mixed-bed IEF configurations; Variability in GFP proteoform profiles demonstrated by intensity plots of GFP focused in µIPG device from MCF7-GFP cell lysate (40,000 cells); Numerical modeling of Nile Blue acrylamide diffusion; Recipes of PA gel precursors; Incubation buffers for all experiments; Anode inlet and cathode inlet sample components for all isoelectric focusing experiments (PDF)

## Author Contributions

All authors designed the experiments. G.L. performed the experiments. G.L. performed the data analysis. The manuscript was written through contributions of all authors. All authors have given approval to the final version of the manuscript.

## ACKNOWLEDGMENT

This work has been supported by the Chan Zuckerberg Biohub Investigator Program (A.E.H), the National Science Foundation Graduate Research Fellowship Program (NSF GRFP) (G.L.), and the Siebel Foundation (G.L.). Photolithography was performed in the QB3 Biomolecular Nanotechnology Center at UC Berkeley. We acknowledge all members of the Herr Lab at UC Berkeley for useful discussions and feedback.

