## Supporting Information for "Reducing Cathodic Drift during Isoelectric Focusing using Microscale Immobilized pH Gradient Gels"

### Supplemental Experimental Section

**µIPG Device Design.** The μIPG device was fabricated by standard soft lithography methods. A silicon wafer (WaferPro C04009) was patterned with SU8 3050 photoresist (MicroChem Y311075) using in-house-designed masks (CAD/ART Services) to create the silicon wafer mold and coated with dichlorodimethylsilane (Sigma 440272) to prevent PDMS from sticking to the SU8 features. RTV615 PDMS (Momentive) was mixed at a 10:1 ratio, degassed for 2 hr under vacuum, and poured over the silicon wafer mold of the microchannel features. Each microchannel is 3.5-mm long, 100-μm wide, and 50-μm tall. After the PDMS was cured at 80 ºC for 2 hr, a biopsy punch was used to make 3-mm diameter inlet and outlet holes for each microchannel. The PDMS slab was cleaned of any debris by sticking and peeling tape to the patterned surface multiple times, and then immediately assembled onto a standard glass slide (VWR 48300-048). Since only light pressure will be applied to the microchannels in subsequent steps, the noncovalent interaction is sufficient to hold the PDMS onto the glass slide during device fabrication and operation. Therefore, we did not perform plasma bonding of the PDMS to glass as is standard in soft lithography. We then apply Kapton tape (Electron Microscopy Sciences 77708-02) on the other side of the glass slide to cover the inlet and outlet holes to serve as a photomask and prevent photopolymerization in those regions.

**Polyacrylamide Gel Fabrication.** To polymerize PA gel in the PDMS channels, we adapted a technique from previous work which ensures gel polymerization in a microchannel environment that may be O_2_ rich^1^. The microchannels are first prepared for PA polymerization by adding 10 μL of a 10% benzophenone (Sigma B9300) in acetone solution to the inlet and outlet and allowing the benzophenone solution to absorb into the PDMS for 3 min. Benzophenone serves as an oxygen scavenger when exposed to UV^2^. The benzophenone solution is then wicked with a Kimwipe and replaced with 10 μL methanol (VWR BDH1135). The methanol is then wicked and a gentle N_2_ stream at one of the inlets is used to dry the microchannels. The methanol rinse and nitrogen drying are repeated one more time. The methanol rinse step removes excess benzophenone solution from the microchannel, as an excess of benzophenone is prone to forming crystals as the acetone evaporates, which could block the microchannel. Next, 10 μL of a 6 %T PA gel precursor made from 30 %T 29:1 acrylamide/bis-acrylamide (Sigma A3574) and UV photoinitiator 2,2-Azobis(2-methyl-N-(2-hydroxyethyl) propionamide) (VA086, Wako Chemicals 01319342) is applied at one of the inlets. Supplemental Table S1 summarizes the PA gel precursor recipes. All PA gel precursors are prepared in deionized (DI) water provided by an Ultrapure Millipore filtration system (18.2 MΩ). Since the benzophenone treatment renders the microchannel hydrophobic, we apply gentle suction at the opposite inlet using a pipette tip to promote the PA gel precursor to fill the microchannel. The scaffold PA gel precursor is then photopolymerized with 2 min of UV light exposure at 20 mW/cm^2^ (measured by an OAI 308 UV intensity meter). An OAI Model 200 Collimated UV light source provides the UV light exposure for gel photopolymerization. The UV activates both the benzophenone (oxygen scavenger) and the VA-086 photoinitiator (radical polymerization initiator).

The inlets were then emptied and filled with 6 %T or 12 %T PA acidic (pH = 3.8) and basic (pH = 7.0) Immobiline-PA gel precursor solutions containing acrylamide/bis-acrylamide, Immobilines, and VA-086 for IPG-IEF or mixed-bed IEF or 6 %T PA gel precursor without Immobilines for CA-IEF (Supplemental Table S1). Acidic Immobilines used were acrylamido buffers pKa 3.6 (∼0.2 M in water, Sigma 01716) and pKa 4.6 (∼0.2 M in water, Sigma 01718). Basic Immobilines used were acrylamido buffers pKa 6.2 (∼0.2 M in 1-propanol, Sigma 01721), pKa 7.0 (∼0.2 M in 1-propanol, Sigma 01729), and pKa 9.3 (∼0.2 M in 1-propanol, Sigma 01738). The acidic and basic IPG precursor recipes were adapted from previous work^3^. The precursors were allowed to diffuse into the microchannels for 7 hr to establish a linear concentration gradient along the length of the microchannel. During this diffusion step, the polymerized 6 %T PA gel serves as a scaffold for the IPG gel, maintaining quiescent conditions during gel precursor diffusion [cite Tentori Microchamber]. The IPG precursor was then photopolymerized with 2 min of UV light exposure at 20 mW/cm^2^. The Kapton tape photomask is then removed from the bottom of the glass slide. The devices were incubated with (for CA-IEF or mixed-bed IEF) or without (for IPG-IEF) 1% ZOOM Carrier Ampholytes pH 4 – 7 (Thermo Fischer Scientific ZM0022) dissolved in DI water or sample loading buffer overnight (as summarized in Supplemental Table S2) and could be stored at 4 ºC for at least 9 days. The sample loading buffer consisted of 3% 3-[(3- Cholamidopropyl)dimethylammonio]-1-propanesulfonate (CHAPS, Sigma RES1300C), 10% D-sorbitol (Sigma S6021), and 200 mM nondetergent sulfobetaine-256 (Abcam ab142233) in DI water.

**pH Gradient Characterization.** To determine the amount of time necessary to establish a linear Immobiline concentration gradient within the microchannel, we monitored the diffusion of Nile Blue acrylamide (Polysciences 25395). Nile Blue acrylamide was spiked into the acidic gel precursor at a concentration of 17.4 μM.

**Polymerization Experiments.** For the polymerization test experiment in Supplemental Figure S1A, Nile Blue acrylamide was spiked into the gel precursors at a concentration of 17.4 μM. For the polymerization test experiment in Supplemental Figure S1B, AlexaFluor-647-labeled donkey anti-rabbit antibody (Invitrogen A-31573) was spiked into the scaffold, acidic, and basic gel precursors at a concentration of 0.01 mg/mL. For the polymerization experiment, the antibody simply served as a large molecular mass dye.

**Cell Culture and Cell Lysate Preparation.** An MCF7 human breast cancer cell line genetically modified to stably express enhanced green fluorescent protein (GFP) was obtained from the American Type Culture Collection, authenticated by short tandem repeat analysis, and tested negative for mycoplasma. The MCF7-GFP cells were maintained in a humidified 37 °C incubator kept at 5% CO2 with RPMI media (Gibco 11875-093) supplemented with 1% penicillin/streptomycin (Invitrogen 15140122) and 10% Fetal Bovine Serum (FBS, Gemini Bio-Products 100-106).

A T75 flask with MCF7-GFP cells at 80% confluency was used to prepare cell lysate. Cell lysate preparation was performed as previously described^4^ with some modifications. Cells were detached with 0.05% Trypsin-EDTA (ThermoFisher 25300-120), resuspended in 4 °C 1× phosphate-buffered saline (PBS, Thermo Fisher Scientific 10010023), and counted with a phase counting chamber (Hausser Scientific 3200). The cell suspension was pelleted, the PBS was removed, and the cell pellet was resuspended in HNTG buffer for a final concentration of 20,000 cells/μL HNTG buffer. HNTG buffer was prepared with 20 mM HEPES pH 7.5 (Sigma H-9897), 25 mM NaCl (Fisher S271), 0.1% Triton X-100 (Sigma X100), 10% glycerol (Sigma G7893), and 1X cOmplete™ EDTA-free Protease Inhibitor Cocktail (Sigma COEDTAF-RO) in DI water. The cells were allowed to lyse in the HNTG buffer for 30 min on ice with vortexing every 5 min. Finally, the samples were clarified by centrifugation at 16,000 × g for 10 min at 4 °C, the lysate supernatant was collected, and 50 μL aliquots were stored at -80 °C.

**Isoelectric Focusing Experiments.** Fluorescent pI markers (pI 4.5, Sigma 89149; pI 5.5, Sigma 77866; pI 5.9, Sigma 89478; pI 6.6, Sigma 73376; pI 6.7, Sigma 73938) were used at various concentrations to provide similar intensities and were diluted in DI water or sample loading buffer. We also tested fluorescent pI marker 4.0 (Sigma 89827), but pI marker 4.0 was excluded from analysis due to difficulty accurately identifying a peak center (data not shown). For all experiments, the anode inlet was filled with 1× IEF Anode Buffer (Bio-Rad 1610761) and the cathode inlet was filled with 1× IEF Cathode Buffer (Bio-Rad 1610762). For cell lysate experiments, 1 μL of MCF7-GFP cell lysate (∼20,000 cells/µL) was applied to both the anode and cathode inlets (~40,000 cells total), as well as pI markers in sample loading buffer. Supplemental Table S3 lists the anode inlet and cathode inlet sample components for all IEF experiments. The inlets were connected to a programmable high voltage power supply, LabSmith HVS448LC 3000V High Voltage Sequencer, with platinum electrodes. An electric field was applied using the following voltage ramp: 50 V/cm for 4 min, 100 V/cm for 5 min, 200 V/cm for 5 min, and 300 V/cm for 6 min or more depending on the experiment.

**Fluorescence Imaging.** The Nile Blue acrylamide diffusion experiment was imaged using an Olympus IX71 inverted epifluorescence microscope equipped with an Andor iXon + EMCCD camera, ASI motorized stage, and X-cite exacte illumination system from Lumen Dynamics. All IEF experiments were imaged using an Olympus IX-51 inverted epifluorescence microscope equipped with a Peltier-cooled CCD camera CoolSNAP HQ2 (Roper Scientific), ASI motorized stage, and X-cite exacte illumination system from Lumen Dynamics. The polymerization experiments (Supplemental Figure S1) were imaged on a GenePix 4300A microarray scanner (Molecular Devices).

**Image/Micrograph Analysis.** Fluorescence micrographs were analyzed using in-house MATLAB (R2022b, MathWorks) scripts modified from previous work^5^. Illumination correction for fluorescence micrographs was performed with the BaSiC plugin for ImageJ 1.54f (NIH). Intensity profiles were generated by summing across the width of the microchannel. Peak center locations and area under the curve (AUC) were calculated by curve-fitting the pI marker peaks from the intensity profiles to a Gaussian function.

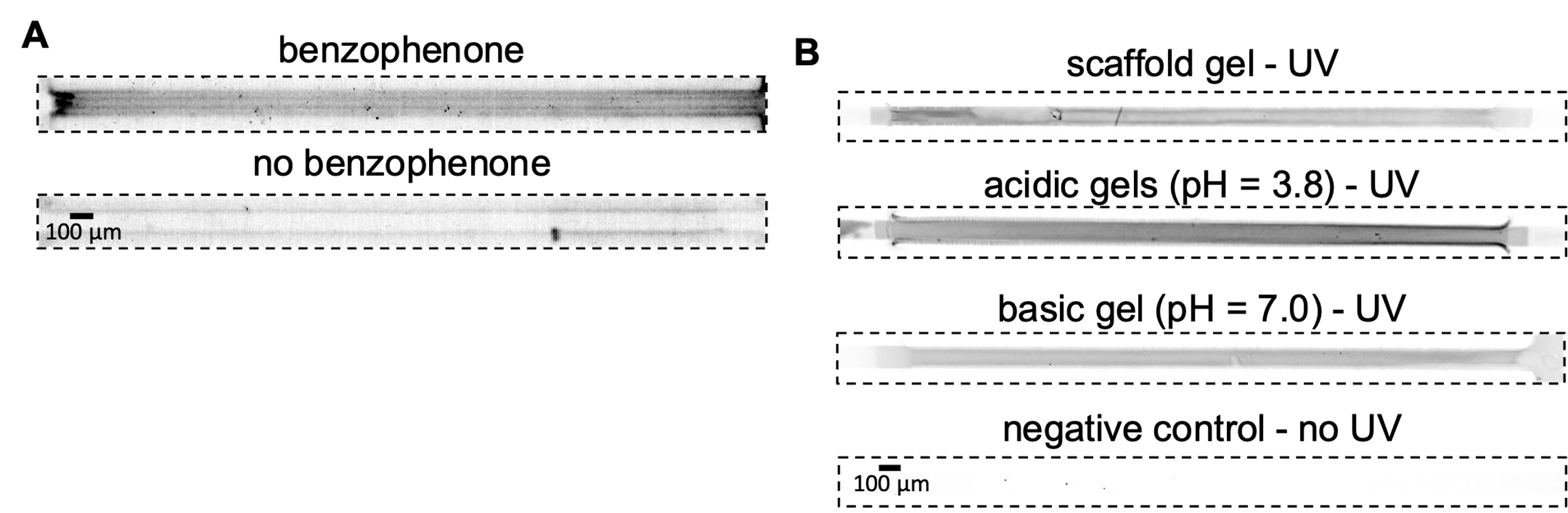

Supplemental Figure S1. Both benzophenone coating and UV irradiation are necessary for PA gel polymerization in the PDMS device. (A) PA gel did not polymerize without benzophenone coating (n = 2 gels). Precursors were spiked with 17.4 µM Nile Blue acrylamide to make the PA gel fluorescent. (B) Confirmation of UV photopolymerization of scaffold, acidic, and basic PA gel precursors individually in microchannels (n = 1 gel). Precursors were spiked with 0.01 mg/mL AlexaFluor-647-labeled antibody to make the PA gel fluorescent. (A-B) Micrographs have the same acquisition settings, brightness, and contrast within panels.

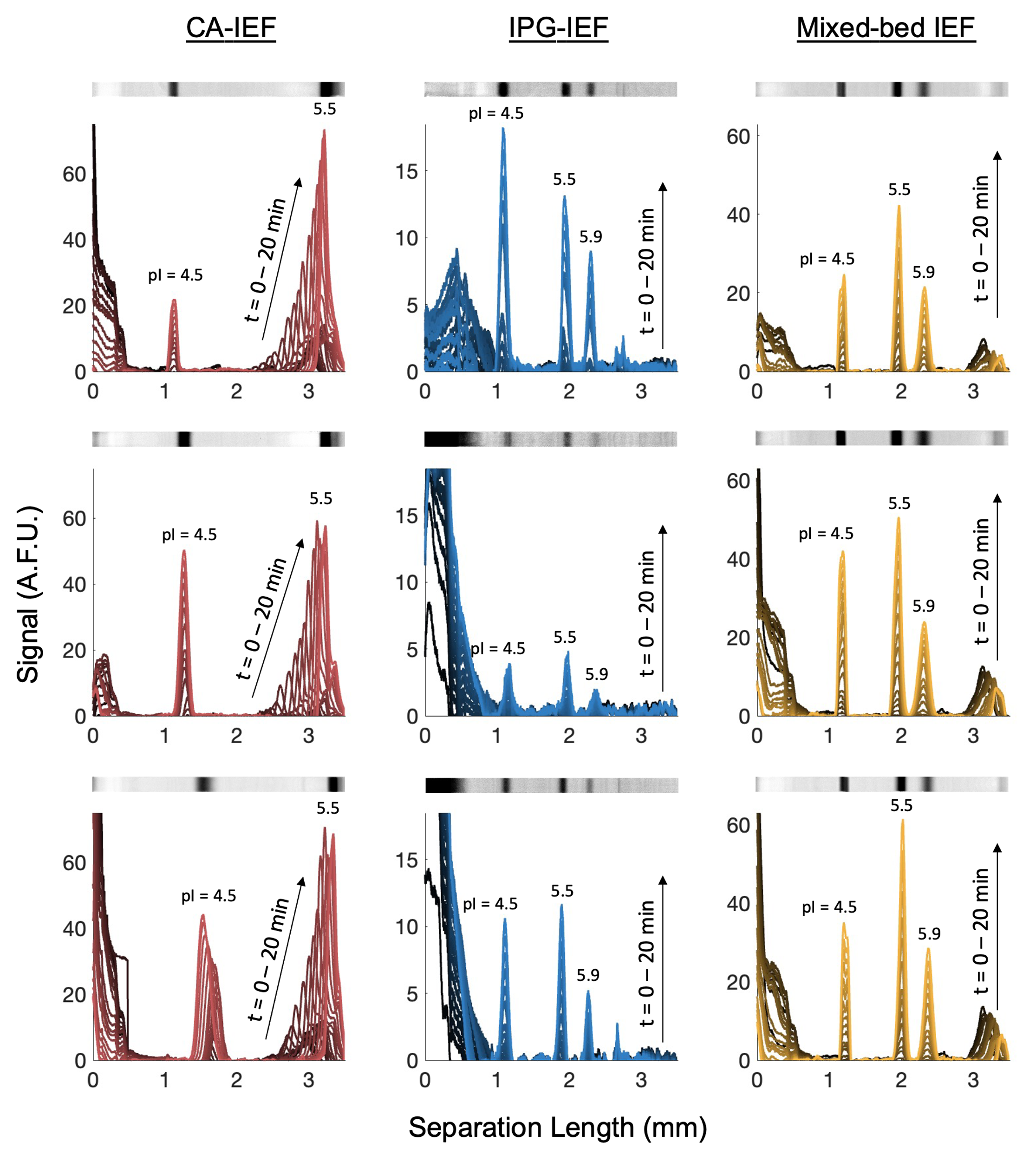

Supplemental Figure S2. Intensity plots and micrographs of pI markers 4.5, 5.5, and 5.9 focused in microfluidic IEF devices in CA-IEF, IPG-IEF, and mixed-bed IEF configurations. Intensity plots are from images taken every minute for 20 minutes of IEF and micrographs above intensity plots are from the final 20-minute time point (n = 3 separations for each IEF configuration). Cathodic drift pf pI marker 5.5 is present in all CA-IEF conditions and anodic drift of pI marker 4.5 is present in the bottom CA-IEF condition.

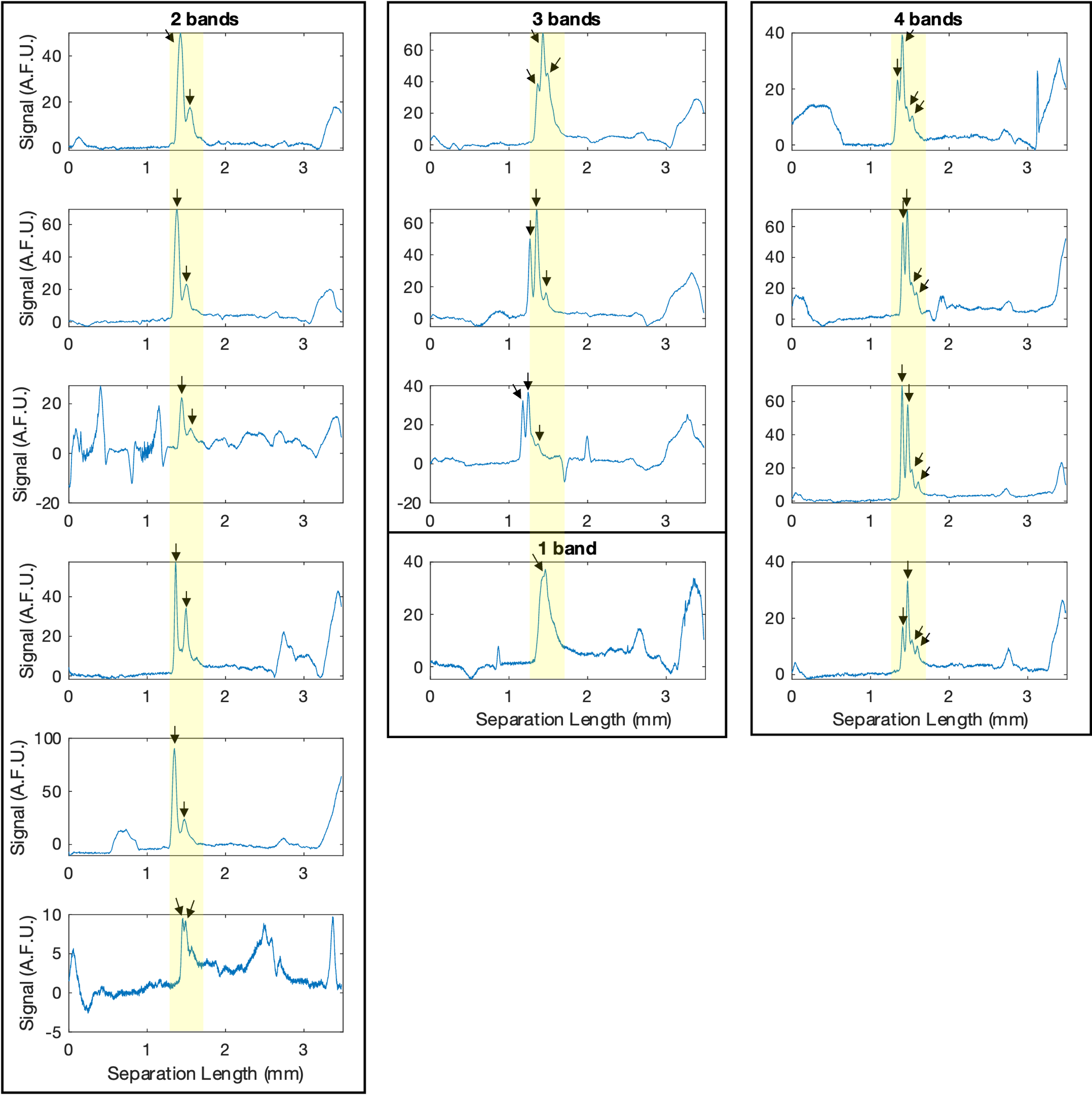

Supplemental Figure S3. Variability in GFP proteoform profiles demonstrated by intensity plots of GFP focused in µIPG device from MCF7-GFP cell lysate (40,000 cells). GFP proteoform peaks are indicated with black arrows. Yellow highlighted bands show consistent positioning of GFP proteoforms within the µIPG microchannel. Profiles captured after 20 min of IEF. IPG gel is 12 %T. IEF performed using mixed-bed IEF.

### Supplemental Note S1. Numerical modeling of Nile Blue acrylamide diffusion.

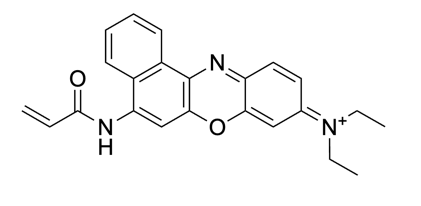

Chemical structure of Nile Blue acrylamide from manufacturer (Polysciences Inc.).

To calculate the diffusion coefficient of Nile Blue acrylamide in PA gel, we used the universal equation for probe diffusion in PA gel^6^ (eq 1) where *D* is the diffusion coefficient of the molecule in the gel (m^2^/s), *D_0_* is the diffusion coefficient of the molecule in the pure solvent (m^2^/s), *R_h_* is the hydrodynamic radius of the molecule (Å), and *C* is the acrylamide concentration (g/mL).

$$\frac{D}{D_{0}}=\exp\left( -3.03R_{h}^{0.59}C^{0.94} \right) (1)$$

To calculate *D_0_* and *R_h_* for Nile Blue acrylamide and solve for *D*, we used the following equations (eq 2, eq 3, and eq 4), which are based on the Stokes-Einstein equation but allow *D_0_* to be estimated from molecular weight^7^. In eq 2, eq 3, and eq 4, *k_B_* is the Boltzmann constant (1.38 × 10^-23^ m^2^kg/s^2^K), *T* is temperature (K), *η* is viscosity of the solvent (kg/ms), *MW* is the molecular weight of the molecule (g/mol), *MW_s_* is the molecular weight of the solvent (g/mol), *ρ_eff_* is the effective density of the solvent (g/m^3^), *N_A_* is the Avogadro number (6.022 × 10^23^ mol^-1^), and *Pa* is the packing fraction of the material (0.64 if a liquid).

$$D_{0}=\frac{k_{B}T\left( \frac{3\alpha}{2}+\frac{1}{1+\alpha} \right)}{6\pi\eta\sqrt[3]{\frac{3MW}{4\pi\rho_{eff}N_{A}}}} \left( 2 \right)$$

$$\alpha=\sqrt[3]{\frac{{MW}_{s}}{MW}} (3)$$

$$R_{h}=\sqrt[3]{\frac{3PaMW}{4\pi\rho_{eff}N_{A}}} (4)$$

If we solve for Nile Blue acrylamide diffusion in a 6 %T PA gel, where water is the solvent, we arrive at *D* = 2.7 × 10^-10^ m^2^/s or 0.0162 mm^2^/min.

Next, the concentration profile of Nile Blue acrylamide in the microchannel was modeled with the 1-D diffusion equation:

$$\frac{\partial c}{\partial t}=D\frac{\partial^{2}c}{\partial x^{2}} \left( 5 \right)$$

where *D* is the diffusion coefficient of Nile Blue acrylamide (0.0162 mm^2^/min), *x* is the spatial coordinate in millimeters, *t* is time in min, and *c* is the Nile Blue acrylamide concentration in micromolar. The initial condition is:

$$c\left( x,0 \right)=0, 0<x<L \left( 6 \right)$$

where *L* is the length of the microchannel (3.5 mm). The boundary conditions are:

$$c\left( 0,t \right)=0, c\left( L,t \right)=U, t>0 \left( 7 \right)$$

where *U* is the concentration of Nile Blue acrylamide in the reservoir (17.4 μM). Applying the technique of separation of variables leads to the analytical solution^8^:

$$c\left( x,t \right)=U\frac{x}{L}-\frac{2U}{\pi}\sum_{n=1}^{\infty} \frac{1}{n}\exp\left( -n^{2}\pi^{2}\frac{Dt}{L^{2}} \right)sin[n\pi\left( 1-\frac{x}{L} \right)] \left( 8 \right)$$

Equation 8 was evaluated every minute from t = 0 to 500 min. Finally, the linearity (R^2^) of the resulting concentration profile at each time point was used to determine the gradient linearity (Figure 2B).

### Supplemental Table S1. Recipes of PA gel precursors.

| **Figure** | **Gel Type** | **Gel Components** |
| --- | --- | --- |
| **1C, 2B, 3, 4, 5, 6, S1, S2, S3** | 6 %T scaffold gel precursor | • 6 %T, 3.3 %C acrylamide/bis-acrylamide  • 0.5 %VA-086 |
| **1C, 2B, 3, 4, 5, S1, S2** | 6 %T pH 3.8 gel precursor | • 6 %T, 3.3 %C acrylamide/bis-acrylamide  • 0.5 %VA-086  • 12.7 mM pKa 3.6 Immobiline  • 7.48 mM pKa 6.2 Immobiline |
| **6, S3** | 12 %T pH 3.8 gel precursor | • 12 %T, 3.3 %C acrylamide/bis-acrylamide  • 1 %VA-086  • 12.7 mM pKa 3.6 Immobiline  • 7.48 mM pKa 6.2 Immobiline |
| **1C, 2B, 3, 4, 5, S1, S2** | 6 %T pH 7.0 gel precursor | • 6 %T, 3.3 %C acrylamide/bis-acrylamide  • 0.5 %VA-086  • 4.40 mM pKa 3.6 Immobiline  • 10.3 mM pKa 4.6 Immobiline  • 2.36 mM pKa 6.2 Immobiline  • 4.17 mM pKa 7.0 Immobiline  • 12.1 mM pKa 9.3 Immobiline |
| **6, S3** | 12 %T pH 7.0 gel precursor | • 12 %T, 3.3 %C acrylamide/bis-acrylamide  • 1 %VA-086  • 4.40 mM pKa 3.6 Immobiline  • 10.3 mM pKa 4.6 Immobiline  • 2.36 mM pKa 6.2 Immobiline  • 4.17 mM pKa 7.0 Immobiline  • 12.1 mM pKa 9.3 Immobiline |

### Supplemental Table S2. Incubation buffers for all experiments.

| **Figure** | **Incubation buffer** |
| --- | --- |
| **1C** | 1% ZOOM Carrier Ampholytes pH 4 – 7 dissolved in sample loading buffer |
| **2B, 3, S1** | DI water |
| **4, 5, S2** | CA-IEF or mixed-bed IEF: 1% ZOOM Carrier Ampholytes pH 4 – 7 dissolved in DI water  IPG-IEF: DI water |
| **6, S3** | Mixed-bed IEF: 1% ZOOM Carrier Ampholytes pH 4 – 7 dissolved in sample loading buffer  IPG-IEF: sample loading buffer |

### Supplemental Table S3. Anode inlet and cathode inlet sample components for all isoelectric focusing experiments.

| **Figure** | **IPG**  **%T** | **Anode Inlet Sample Components** | **Cathode Inlet Sample Components** |
| --- | --- | --- | --- |
| **1C** | 6 | • 7.5 *µ*L of pI 4.5 marker (10 *µ*g/mL), pI 5.5 marker (30 *µ*g/mL), and pI 5.9 marker (20 *µ*g/mL) in sample loading buffer  • 1 *µ*L cell lysate (*∼*20,000 cells)  • 6.5 *µ*L DI water  • 1.5 *µ*L Anode Buffer | • 7.5 *µ*L sample  loading buffer  • 1 *µ*L cell lysate  (*∼*20,000 cells)  • 6.5 *µ*L DI water  • 1.5 *µ*L Cathode  Buffer |
| **3** | 6 | • 5 *µ*L of pI 4.5 marker (10 *µ*g/mL), pI 5.5 marker (30 *µ*g/mL), pI 5.9 marker (20 *µ*g/mL), pI 6.6 marker (20 *µ*g/mL), and pI 6.7 marker (20 *µ*g/mL) in DI water  • 10 *µ*L DI water  • 1.5 *µ*L Anode Buffer | • 15 *µ*L DI water  • 1.5 *µ*L Cathode  Buffer |
| **4, 5, S2** | 6 | • 7.5 *µ*L of pI 4.5 marker (10 *µ*g/mL), pI 5.5 marker (30 *µ*g/mL), pI 5.9 marker (20 *µ*g/mL), in DI water  • 7.5 *µ*L DI water  • 1.5 *µ*L Anode Buffer | • 15 *µ*L DI water  • 1.5 *µ*L Cathode  Buffer |
| **6, S3** | 12 | • 7.5 *µ*L of pI 4.5 marker (10 *µ*g/mL), pI 5.5 marker (30 *µ*g/mL), and pI 5.9 marker (20 *µ*g/mL) in sample loading buffer  • 1 *µ*L cell lysate (*∼*20,000 cells)  • 6.5 *µ*L DI water  • 1.5 *µ*L Anode Buffer | • 7.5 *µ*L sample  loading buffer  • 1 *µ*L cell lysate  (*∼*20,000 cells)  • 6.5 *µ*L DI water  • 1.5 *µ*L Cathode  Buffer |
